# MatchMiner: An open source computational platform for real-time matching of cancer patients to precision medicine clinical trials using genomic and clinical criteria

**DOI:** 10.1101/199489

**Authors:** James Lindsay, Catherine Del Vecchio Fitz, Zachary Zwiesler, Priti Kumari, Bernd Van Der Veen, Tamba Monrose, Tali Mazor, Susan Barry, Adem Albayrak, Michael Tung, Khanh Do, Suzanne Hector-Barry, Brian Beardslee, Geoffrey Shapiro, John Methot, Lynette Sholl, Laura E. MacConaill, Neil Lindeman, Bruce Johnson, Barrett Rollins, Chris Sander, Michael Hassett, Ethan Cerami

## Abstract

**Background:** Molecular profiling of cancers is now routine at many cancer centers, and the number of precision cancer medicine clinical trials, which are informed by profiling, is steadily rising. Additionally, these trials are becoming increasingly complex, often having multiple arms and many genomic eligibility criteria. Currently, it is a challenging for physicians to match patients to relevant clinical trials using the patient’s genomic profile, which can lead to missed opportunities. Automated matching against uniformly structured and encoded genomic eligibility criteria is essential to keep pace with the complex landscape of precision medicine clinical trials.

**Results:** To meet these needs, we built and deployed an automated clinical trial matching platform called MatchMiner at the Dana-Farber Cancer Institute (DFCI). The platform has been integrated with Profile, DFCI’s enterprise genomic profiling project, which contains tumor profile data for >20,000 patients, and has been made available to physicians across the Institute. As no current standard exists for encoding clinical trial eligibility criteria, a new language called Clinical Trial Markup Language (CTML) was developed, and over 178 genomically-driven clinical trials were encoded using this language. The platform is open source and freely available for adoption by other institutions.

**Conclusion:** MatchMiner is the first open platform developed to enable computational matching of patient-specific genomic profiles to precision cancer medicine clinical trials. Creating MatchMiner required developing clinical trial eligibility standards to support genome-driven matching and developing intuitive interfaces to support practical use-cases. Given the complexity of tumor profiling and the rapidly changing multi-site nature of genome-driven clinical trials, open source software is the most efficient, scalable, and economical option for matching cancer patients to clinical trials.

## Background

Precision cancer medicine has begun to transform cancer care. Examples are numerous, including the use of tyrosine kinase inhibitors (TKIs) for chronic myelogenous leukemia (CML), epidermal growth factor receptor (EGFR) and anaplastic lymphoma kinase (ALK) inhibitors for lung cancer, BRAF inhibitors for melanoma, and pembrolizumab for microsatellite instability-high advanced cancers [1, 2, 3, 4, 5]. As genome sequencing becomes an integral part of cancer care [6, 7], and the set of new experimental drug targets expands [8, 9], there is an increasingly urgent need to efficiently match patients to genotype-driven clinical trials [10, 11]. In the absence of advanced informatics systems, individual oncologists must track hundreds of active clinical trials, only a few of which may be relevant for any given patient at a time [12, 13]. Likewise, clinical trial investigators are frequently challenged to manually evaluate pools of hundreds or thousands of patients in order to identify the few eligible patients for their specific trial. The challenge is greatly exacerbated across multiple cancer centers, where thousands of trials may be active and each institution may have used different standards for defining clinical trial criteria [9,14]. Further, many precision medicine basket trials aim to enroll patients across multiple histologies, each of which is defined by unique genomic criteria, and the logistics of identifying eligible patients for such trials can be daunting [14, 15, 16].

Few (<5%) cancer patients currently participate in clinical trials [17, 18, 19]. However, there is emerging evidence that advanced automated informatics platforms can greatly improve trial matching and increase participation rates. For example, a recent study from Memorial Sloan Kettering Cancer Center detailed their enterprise sequencing effort coupled with automatic notification of new patient matches and reported an 11% enrollment rate to genotype-matched clinical trials [20]. A similar match rate has been reported at MD Anderson Cancer Center, which has also developed its own trial matching infrastructure [21]. Multiple informatics platforms have been developed to address the specific challenges associated with matching of patients to precision cancer medicine clinical trials. These include academic cancer center-specific solutions [12, 21] and commercial solutions from companies such as Foundation Medicine, IBM Watson, Syapse, Genospace, and GenomOncology. Some of these platforms have already influenced clinical care. However, these solutions are either proprietary or not currently portable to other institutions, severely limiting their wide adoption. This confines automated trial matching to a few select cancer centers and limits the overall rate at which precision cancer medicine clinical trials accrue. There is also no open standard by which genomic eligibility criteria for clinical trials can be encoded uniformly across cancer centers, which further limits the portability of existing systems. An open software platform and a genomic eligibility standard would address these issues and greatly reduce the logistical burden of precision cancer medicine trial recruitment, increase the overall rate of trial accrual, maximize trial options for individual patients, and increase multisite collaboration.

## Implementation

MatchMiner is an open source computational platform for algorithmically matching patients to precision medicine clinical trials based on clinical and genomic criteria (Figure 1). The following are inputs to MatchMiner: 1) patient-specific genomic data, including somatic mutations, single nucleotide variants (SNVs), copy number alterations (CNA), and structural rearrangements; 2) patient-specific clinical data, currently including primary cancer type, gender, and age; 3) structured eligibility criteria for clinical trials, including detailed arm-level inclusion and exclusion criteria; and 4) real-time clinical trial status information, indicating overall trial status and the individual statuses of each arm within a given trial. The MatchMiner platform automatically matches the patient-specific genomic events to clinical trials, and makes the results available to trial investigators and clinicians via a HIPAA compliant web-based platform.

**Figure 1.**
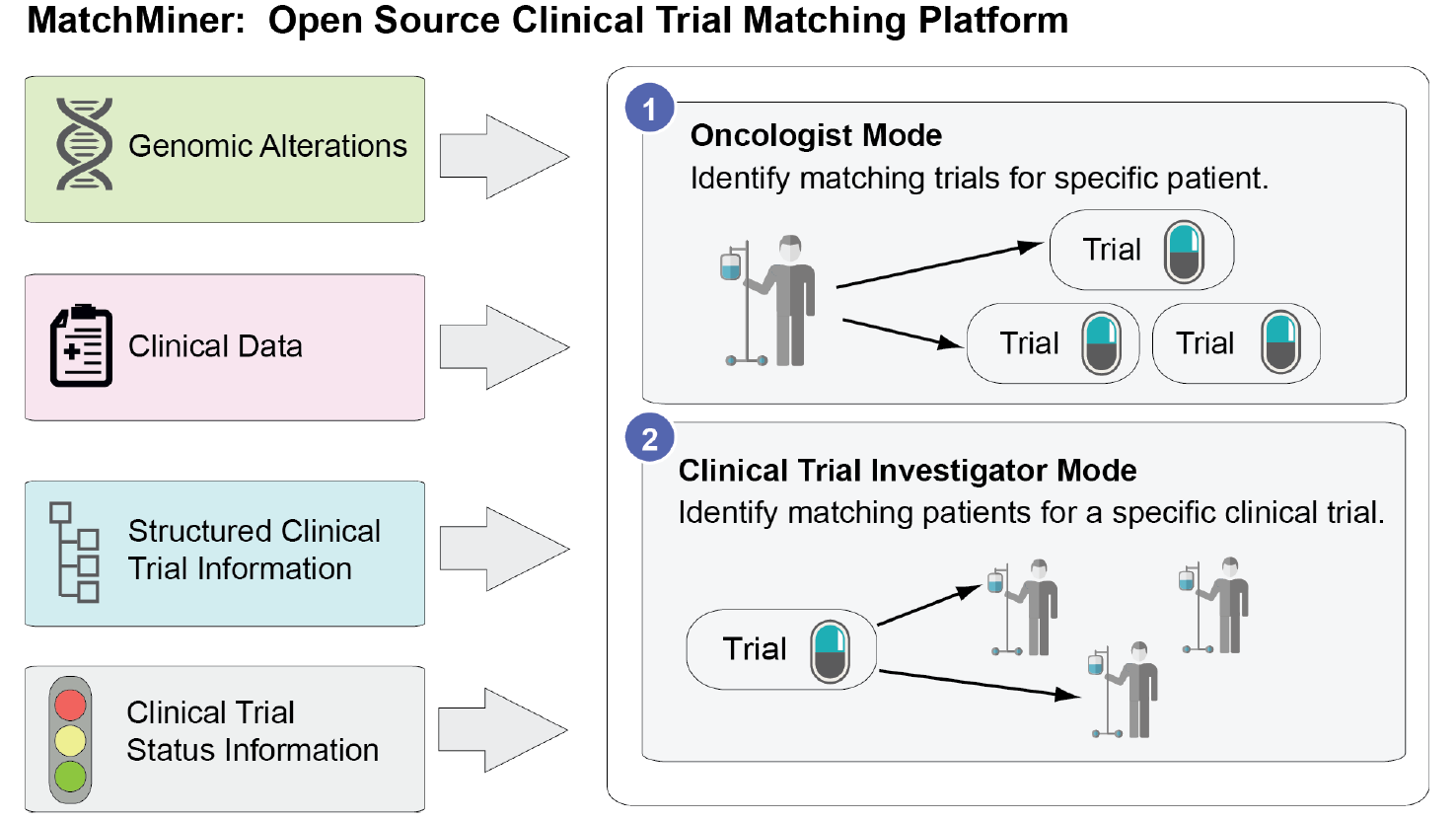
MatchMiner overview. The four inputs to MatchMiner (left) and the two modes of use (right) are illustrated. Oncologist mode allows physicians to quickly review precomputed matches for a particular patient. The clinical trial investigator mode allows investigators to search for patients who have been genetically profiled, and get alerted when new patients are sequenced that matched specific criteria.

**Figure 2.**
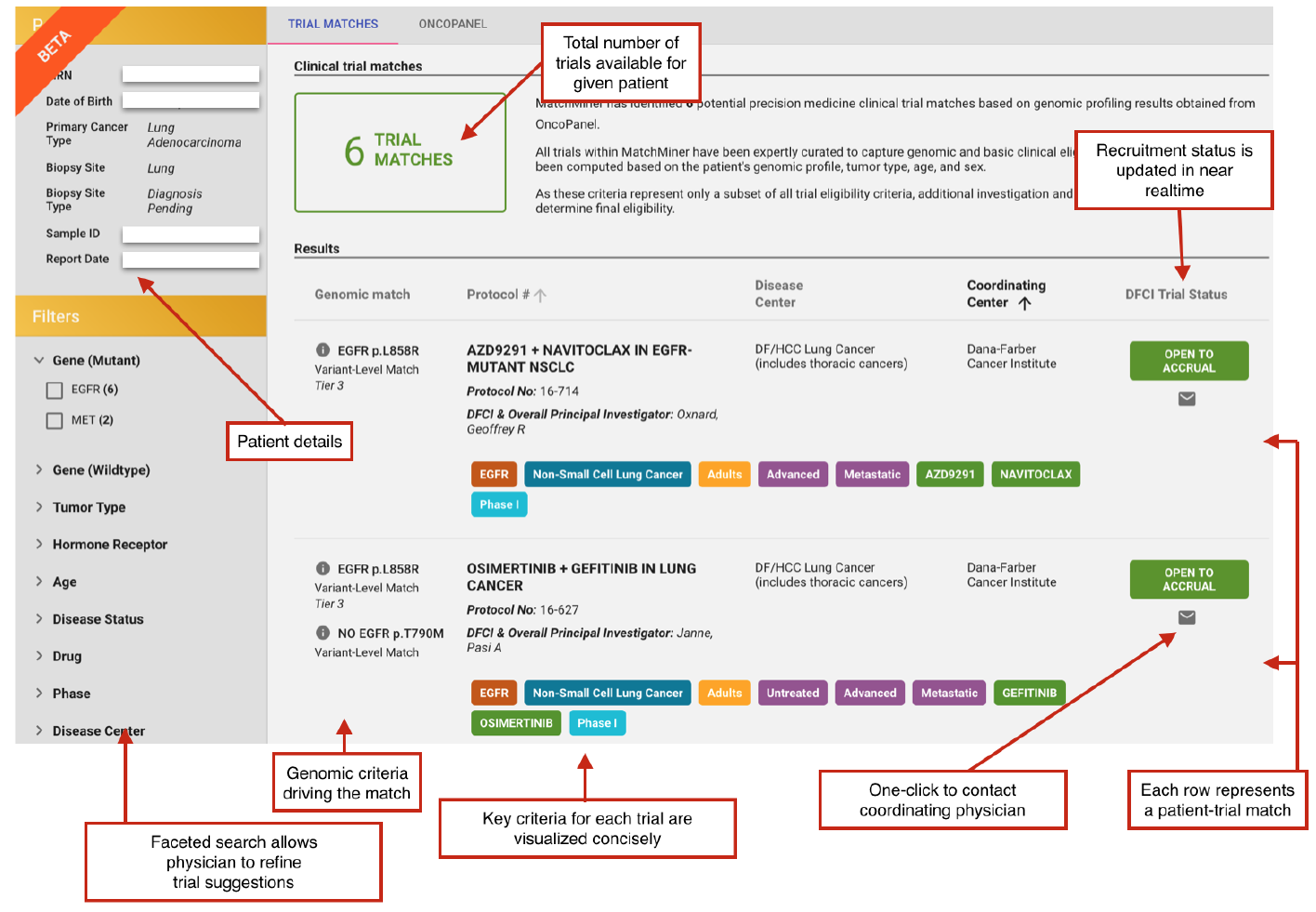
Patient trial matches. Screenshot of MatchMiner in the oncologist mode, showing genomically driven clinical trial matches for a single patient (simulated patient data). Genes specified for inclusion/exclusion, disease, age restrictions, stage restrictions, therapeutics and phase are considered key criteria for a trial.

MatchMiner currently supports two modes of use (Figure 1). In the *oncologist* mode, MatchMiner can enable clinicians to use the genomic data of their patient to retrieve trial matches for that specific patient or find trials using a robust search interface. Upon entering a patient’s name or medical record number, a patient-centric view is displayed that provides a visual summary of trial matches and the patient’s genomic changes generated by the Profile program. When patients have external genomic reports (e.g., from Foundation Medicine instead of DFCI’s Profile program) that are not part of the automated data feed, manual trial search can be performed. Keywords can be manually copied from the external report, and results can be refined using a faceted search. The faceted search interface allows for filtering based on clinical and genomic criteria, including gene, cancer type, trial status, trial phase, etc.

In the *clinical trial investigator* mode, investigators can create a query to identify patients for specific trials based on genomic and clinical criteria, and monitor the patient cohort for new potential trial matches. Investigators are automatically notified via email of new potential matches and are provided with a real-time dashboard of patient matches (Figure 3A). The platform also provides workflow options for reviewing patient matches, and an email templating system to streamline the process of contacting treating physicians. Finally, clinicians can identify aggregate statistics regarding patient matches, including monthly accrual rates, based on historical data (Figure 3B).

**Figure 3.**
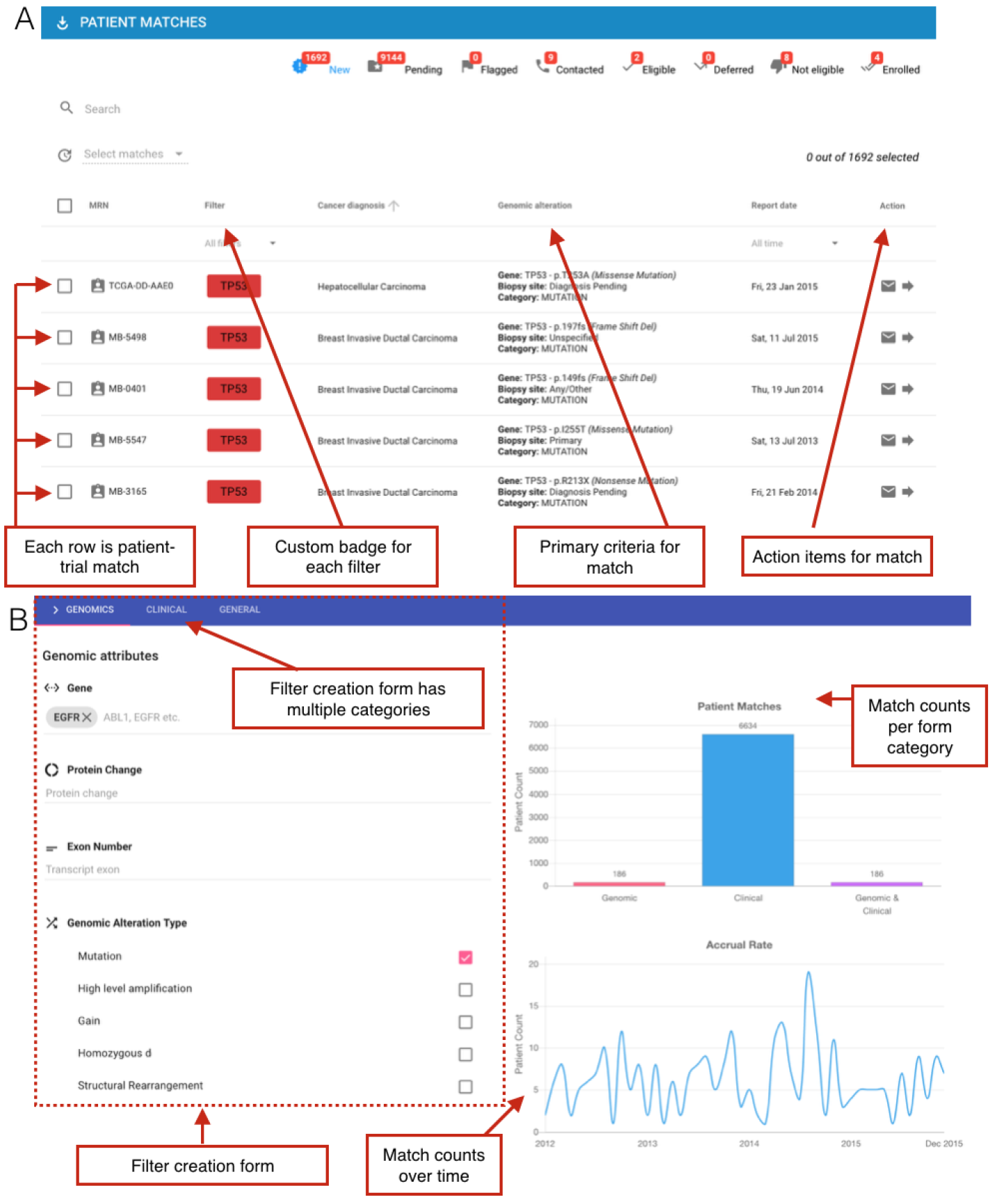
Clinical trial investigator mode. Screenshots of MatchMiner in the clinical trial investigator mode (simulated patient data). A) Dashboard of patient matches for a particular clinical trial. B) Filter creation view for identification of specific patients for a clinical trial, including aggregate statistics for forecasting clinical trial enrollment.

MatchMiner is a two-tier web application with a Python-based REST application programming interface (API) server and AngularJS 1.5 client. MongoDB is used for data persistence. Microservices are used to load genomic and clinical trial data. Additional microservices can be deployed with MatchMiner for connecting to an electronic medical record (EMR) and clinical trial management systems. For example, DFCI utilizes OnCore from Forte Research Systems, a commonly used system for managing clinical trials within large academic medical centers. Authentication of users is performed via Security Assertion Markup Language (SAML), enabling MatchMiner to be integrated with existing authentication and security systems. When hosted within a secure institutional firewall, MatchMiner is fully HIPAA-compliant, and automatically logs all data requests that expose protected health information. MatchMiner software is open source and available under a GNU Affero License. Source code is available at http://matchminer.org and https://github.com/dfci/matchminer.

Patient genomic and clinical data are stored in a database in two collections. Matchable criteria encoded in CTML are translated into two database queries, one for the clinical criteria and one for the genomic criteria. CTML requires the matchable criteria to be expressed as a Boolean logic clause composed of “and”, “or”, and “not” statements. The logical hierarchy forms a Boolean binary expression tree which is used to programmatically generate database queries. Internal nodes of the tree can be “and” nodes where the intersection of the constituent patient sets is taken, or “or” nodes where the union operation is used. The cumulative patient set for the clauses is computed in a bottom-up manner. For a given trial, both the genomic and clinical queries return a patient set, and the matches are the intersection of the patients in the two sets.

## Clinical trial markup language

CTML is a human-readable programming language that we developed for describing clinical trial eligibility criteria. Clinical trial protocols are large formal documents which specify many aspects of trials, and eligibility criteria is communicated in a variety of forms, including free text, lists, and tables. Concepts including mutation or CNA are not always formally defined within a protocol and often are not standardized across protocols. CTML provides a mechanism to structure that information. CTML currently supports genomic, clinical, and demographic criteria, and can be expanded to encompass any desired eligibility criteria.

CTML files are structured similarly to clinical trials in that they have an intrinsic tree-like structure that is anchored by the core trial details and then extended by arms and dose levels. Each arm and dose level component of a trial may have its own distinct eligibility criteria, which can be encoded in the CTML document.

CTML has five components:

- **Trial**: The core component which contains basic metadata, such as its short and long titles, the national clinical trial (NCT) purpose and identifier from the public registry of trials at clinicaltrials.gov, contact information, and phase of the study.
- **Treatment**: This component describes the steps, arms, doses, and expansion cohorts of a clinical trial, which can form a tree-like structure for complex trials. Each node of this tree can have match criteria which contains genomic and clinical eligibility information.
- **Match**: Contains Boolean logic specifying genomic and clinical criteria. Additional match components can be referenced via logical “and/or/not” operators.
- **Clinical**: A Boolean clause that contains specific clinical criteria. Currently supported fields are age, gender, and histology.
- **Genomic**: A Boolean clause that contains specific genomic criteria. Currently supported fields are Human Genome Organization (HUGO) gene symbol, wild type flag, specific exons, specific variants, specific variant categories, gene level copy number call, and structural variants.

CTML was designed to be easily machine readable and human writable. In general, the relevant clinical and genomic information are pulled from the unstructured free text and converted into a logical expression inside the match clauses. One critical component to accurately matching is ensuring consistent disease labels between the trial definitions and patient medical record data. To address this DFCI adopted the cBioPortal OncoTree ontology, a cancer type taxonomy now used by Memorial Sloan Kettering Cancer Center and other members of the American Association for Cancer Research (AACR) GENIE consortium [22]. Figure 4 shows the CTML encoding of supported fields for an example clinical trial. At the root of the CTML document is basic metadata describing the trial. The structure of the trial is encoded in the Treatment List field. Two arms are shown: one that has two qualifying mutations in BRAF and the other that requires wild-type BRAF. Both arms require the patient to be older than 18 years of age and have a melanoma tumor type that is exclusive of uveal, ocular, and mucosal melanoma.

**Figure 4.**
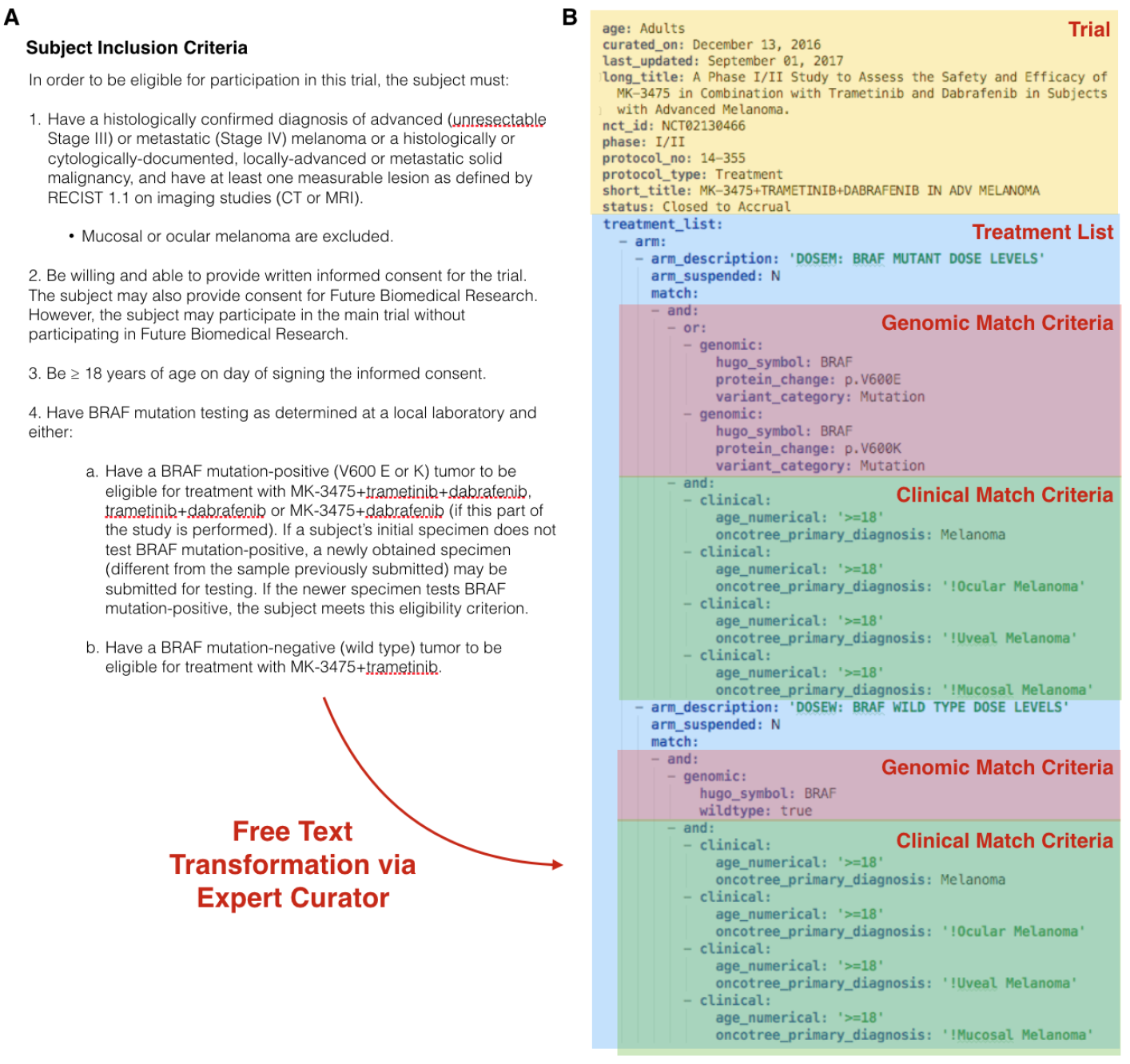
Clinical trial markup language (CTML) Example of clinical trial NCT02130466 encoded in the Clinical Trial Markup Language. A) Excerpt of inclusion criteria represented in free text format. To enroll in this trial, patients must have a histologically confirmed melanoma, excluding ocular, uveal, and mucosal melanomas, and have either wild-type BRAF or a documented BRAF V600E or V600K mutation (for different arms of the trial). B) Trial details transformed into CTML, with curated information related to basic metadata (yellow) and the treatment arms (blue) containing specific genomic and clinical (green) eligibility criteria.

## Results

MatchMiner is now available within DFCI and is fully integrated into Profile, DFCI’s enterprise sequencing effort available to all cancer patients at DFCI, Brigham and Women’s Hospital, and Boston Children’s Hospital[23]. The Profile project uses MPS to identify mutations, CNAs, and structural variations in approximately 450 cancer relevant genes. All genomic events are reviewed by molecular pathologists at Brigham and Women’s Center for Advanced Molecular Diagnostics (CAMD), and mutations are tiered on a scale of 1 to 4 according to clinical actionability [23], and alterations of unknown significance are categorized as tier 4.

As part of our DFCI roll-out, the IRB approved protocol governing the Profile project was amended to include guidelines regarding institutional use of MatchMiner. Enrollment in the Profile project uses a universal consent process, and only patients who specifically agree to use the genomic results from Profile in their medical care are matched within the MatchMiner platform. Use of MatchMiner is restricted to clinical trial enrollment only, not general research use, and only clinical trial investigators with active trials and authorization by the DFCI Chief Clinical Research Officer can access the trial-centric matching. Currently, 708 physicians and approved clinical trial support staff have access to the patient-centric version of MatchMiner, and 129 of these users also have trial-centric access. Over the first 5 months of availability, there have been 1,594 visits to the site by 80 independent investigators, with an average 4.5 minutes of use. A study of MatchMiner’s impact, including plans to record trial enrollments is detailed in the discussion section.

Genomic data generated by the Profile project is automatically transferred to MatchMiner on a nightly basis, with a full suite of automated data quality checks run prior to import. The quality checks include testing for completeness of the data transfer, and ensuring that all clinical and genomic information follows specified standards. The platform also regularly imports data nightly from various DFCI-specific information platforms, including patient-consent information from the consented research database, clinical data from the EHR, and trial status information from the central DFCI clinical trial protocol and enrollment platform, currently running in OnCore. The software platform itself runs within a secure data facility and firewall, and it meets all security and HIPAA requirements defined and audited by the hospital Information Systems team. Clinical trials are curated centrally by a Ph.D.-level biologist with extensive experience in clinical genomics. Any trial open at DFCI with genomic criteria that can be assessed via the Profile test is used for matching within MatchMiner, and new trials are incorporated in a weekly review of protocols. Thus far, MatchMiner currently excludes trials with solely gene expression criteria and other biomarkers that cannot be detected by DNA MPS in a CLIA lab.

There are currently 600 open interventional clinical trials at DFCI, of which 178 are genotype-driven trials. We have curated these trials and they are available in MatchMiner. Curated trials have genomic eligibility criteria ranging from simple, requiring only a single genomic variant criterion (43%), to complex multiple variant criteria with 6 or more genomic changes as seen in Figure 5A. Almost half of trials specify a single cancer type, over 19% of trials list multiple cancer types, and trials targeting all solid or all liquid histologies make up the remaining 32% (Figure 5B). A total of 136 genes are used as eligibility criteria at DFCI (Supplementary Table S1). This list subsumes the 38 trial eligibility genes reported in a survey of precision medicine trials listed on clinicaltrials.gov [9], which indicates the rapidly expanding use of biomarkers and genomic changes as trial eligibility. The genes most commonly required by trials at DFCI include EGFR (18%), ALK (15%), KRAS (15%), and BRAF (15%) (Supplementary Table S1). These are well-studied oncogenes that are often required by clinical trials at other institutions as well [9].

**Figure 5.**
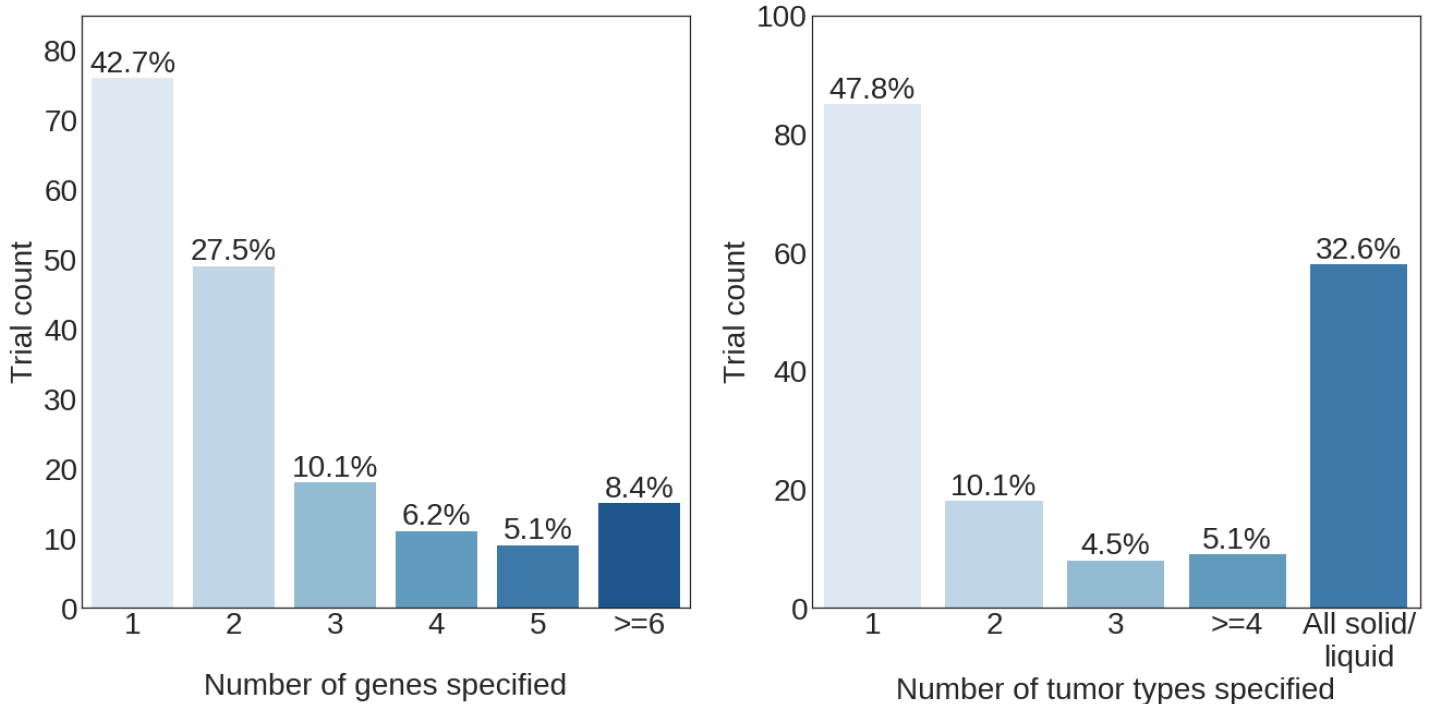
Trial complexity. Analysis of the number of genes (A) and tumor types (B) specified in eligibility criteria for each of the curated trials at DFCI. Trials that specify all solid or all liquid tumors in any arm are only counted in that category, even if other specific tumor types are mentioned.

The most frequently specified genes in trial eligibility criteria are found in multiple cancer types (Figure 6). Many genes which are well-established biomarkers for a given therapy in one disease, e.g. EGFR in non-small cell lung cancer [24, 25], are also being evaluated as biomarkers in other diseases through basket or all solid tumor studies [26]. Precision medicine trials targeting all solid tumors are most frequent at DFCI, likely due to the changing design of trials in the era of genomic medicine and the increasing use of basket studies[27, 15].

**Figure 6.**
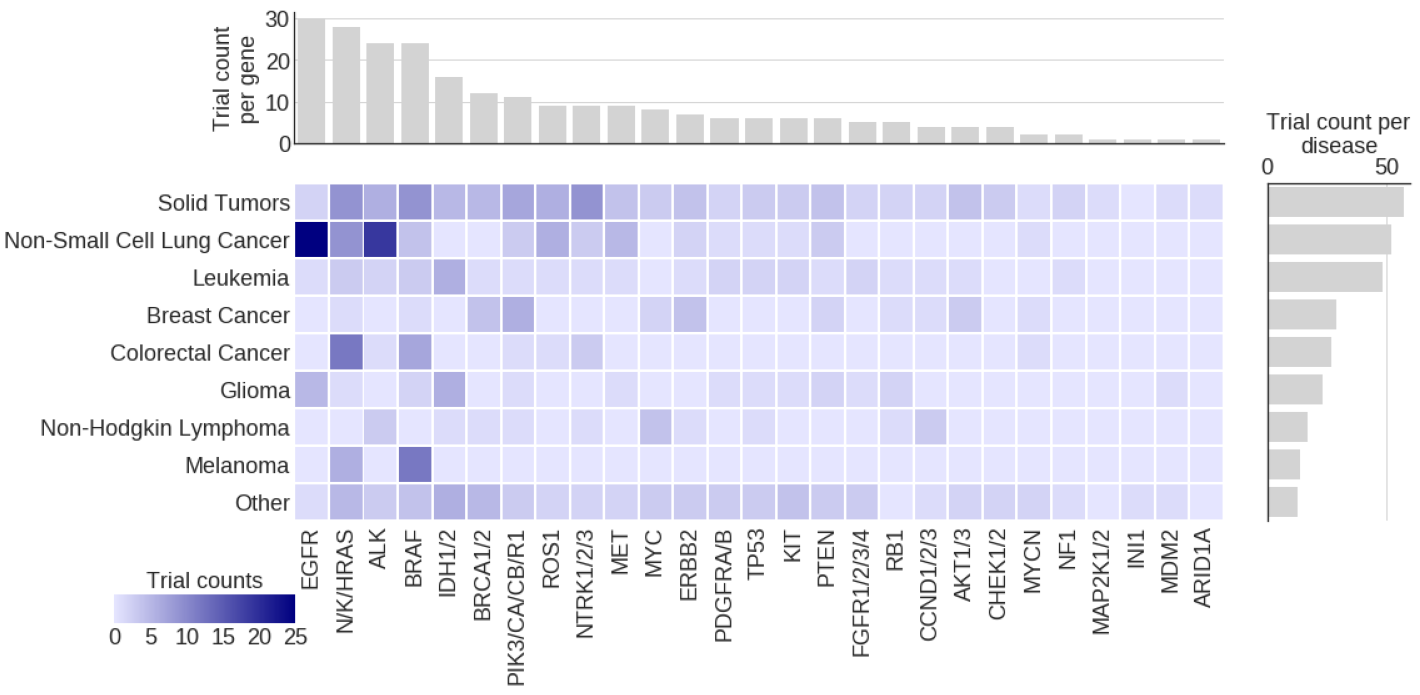
Genomic eligibility summary for clinical trials at DFCI. Overview of the frequency of gene use in eligibility criteria for DFCI trials targeting the top cancer types. In total, 178 trials were included in the analysis. A cell in the heatmap represents the number of trials in which the gene and cancer type were found together. If multiple genes and/or cancer types are specified, all valid combinations are counted, but only once per trial. The bar plot on the top indicates the total number of trials in which the given gene or gene family are mentioned. The bar plot to the right conveys the number of trials that target the given cancer type. Only the most 27 frequently specified gene families are included in the plot.

## Discussion

Clinical trials are a cornerstone of the development of safe and effective new therapies for patients with cancer. More than 10,000 interventional studies for cancer are registered on clinicaltrials.gov and are currently open to enrollment; many of these treatments target specific molecular alterations. Tumor profiling tests usually interrogate hundreds of molecular targets and generate vast amounts of data. With thousands of potential trials, hundreds of potential therapeutic targets, and increasingly complex trials, finding the right trial for the right patient has become an even more challenging task. A few proprietary health information technology platforms have attempted to tackle this problem, but these tools have major limitations and none has seen widespread adoption.

To address this critical need, our team at the Dana-Farber Cancer Institute (DFCI) developed MatchMiner - a novel open source computational platform for matching cancer patients to precision medicine clinical trials using patient-specific genomic profiles and clinical criteria. The platform currently contains tumor profile data for >15,000 patients and structured eligibility criteria for >178 precision medicine trials. Since its launch in late 2016, oncologists in the Early Drug Development Center have reported successful enrollments of patients on clinical trials aided by the MatchMiner platform.

Future efforts will provide additional details about the number of patients assigned to trials using the MatchMiner platform and rate of uptake by the clinical investigators and other providers. The quantitative evaluation of usability involves analyzing temporal trends in utilization by different providers and the number of times the providers access the system. Specifically, we are tracking the following metrics on a monthly basis: the number of unique users who access MatchMiner at least once; the mean number of days each user accessed MatchMiner; the average length of a MatchMiner session; where users exited the session (middle versus end of the workflow); the number of match alerts issued; the proportion of patients with tumor profiling results and clinic visits for whom match alerts were issued; the proportion of clinicians for whom match alerts were issued; and the actions taken for match alerts (enrollment or not), and number of patients enrolled in clinical trials following MatchMiner use. We will analyze performance for these metrics relative to the implementation date using run charts, which is a straightforward analytical tool used frequently in quality and patient safety research [28]. We expect improvements in these performance metrics following tighter integration of MatchMiner into the DFCI’s EMR system.

To evaluate the impact of MatchMiner on clinical trial participation, we will conduct a temporal analysis comparing participation rates before and after implementation of the enhanced/EMR-integrated version of MatchMiner. The primary outcome will be the patient incidence rate, calculated as the number of unique patients who enrolled in a clinical trial during the observation period divided by the number of unique patients seen during the observation period. We hypothesize that there will be a significant increase in the patient incidence rate after the implementation date.

As MatchMiner has been deployed within DFCI, a number of challenges have been identified that will inform its future evolution. The first such challenge is integrating MatchMiner into existing clinical workflows. Profile and MatchMiner were initially developed as clinical research platforms, and they remain largely independent of the main clinical workflows in the institutional EMR system. Clinicians are therefore challenged to integrate MatchMiner into their daily clinical workflow. Fully integrating MatchMiner into an EMR system is challenging, especially for an open source platform that aims to be EMR-vendor agnostic. We are exploring open standards for future releases, such as the SMART on FHIR, a set of open specifications to integrate apps with EMRs [29, 30].

A second major challenge is integrating non-genomic eligibility criteria, such as prior therapies and laboratory values, into clinical trial matching. Given the wide array of such criteria, this is a much greater challenge than integrating genomic data. It also requires modifying the Clinical Trial Markup Language (CTML) to include non-genomic criteria, and interoperable standards for extracting structured clinical data from EMRs. We envision proposing a formal standard for CTML via widely used collaborative standard development processes such as the Global Alliance for Genomics and Health (GA4GH).

Lastly, most major academic cancer centers in the U.S. currently implement their own unique next-generation sequencing platform, and each of these platforms is embedded in larger unique institutional systems for consenting patients, annotating cancer variants, governing who has access to such data, and integrating genomics data into an EMR. We have endeavored to isolate the DFCI-specific portions of MatchMiner into specific microservices and configuration files. However, it is likely that other institutions that adopt MatchMiner will also need to build such microservices and further adapt the code to meet their institutional requirements. As we build an open source community around MatchMiner, our hope is that the community will lead the development of best practices and reusable microservices that can be used across cancer centers.

## Conclusion

MatchMiner is an open source computational platform for algorithmically matching patients to clinical trials using patient-specific genomic profiles and clinical criteria. The platform aims to increase accrual in genotype-specific trials, and maximize trial options for individual patients. The platform is being piloted at DFCI, where it has been integrated into Profile, DFCI’s enterprise next-generation sequencing platform that is freely available to all DFCI patients.

The open-source nature of MatchMiner is one of the key differentiators between this effort and other trial matching initiatives. The source code for the platform is open and available to other cancer centers that wish to integrate automated trial matching with their existing genomic profiling efforts. Additionally, CTML is designed to be an open standard that will evolve as it is adopted by more users and as a greater variety of matchable criteria are integrated. Ideally, the eligibility criteria for trials will also be made more transparent. With a focus on collaboration and open standards, we hope to lessen the burdens associated with manual matching of patients to genomically driven clinical trials, increase trial accrual, and improve long-term outcomes for cancer patients.

## Availability and requirements

Project Name: MatchMiner

Project Home Page: https://matchminer.org

Project Repository: https://github.com/dfci/matchminer

Operating System(s): Docker compatible OS, e.g., Linux, Mac, Azure, Windows

Programming Languages: Python, HTML, Javascript

Other requirements: Clinical trial and patient genomic data

License: GNU Affero License

## Abbreviations

AACR: American Association for Cancer Research
API: Application programming interface
CNA: DNA copy number alteration
CLIA: Clinical Laboratory Improvement Amendments
CTML: Clinical Trial Markup Language
DFCI: Dana-Farber Cancer Institute
EMR: Electronic Medical Record
FFPE: Formalin-fixed paraffin-embedded
GA4GH: Global Alliance for Genome and Health
HIPAA: Health Insurance Portability and Accountability Act
HUGO: Human Genome Organization
Indel: Insertion / deletion
NCT: National Clinical Trial
MPS: Massively Parallel Sequencing
REST: RepresEntational State Transfer
SNV: Single Nucleotide Variant
SAML: Security Assertion Markup Language

## Declarations

### Ethics approval and consent to participate

Use of MatchMiner at DFCI is covered by the core institutional protocol for genomic profiling (Profile) governed by IRB protocol 11–104.

### Consent for publication

Not applicable

### Availability of data and material

The majority of the genomic data analysed in this study are available through AACR Project GENIE (http://sagebase.org/research-projects/aacr-project-genie/).

Due to legal restrictions, curated clinical trial eligibility criteria are not available for public release. However, a subset of trial information is available on ClinicalTrials.gov.

### Authors’ contributions

EC conceived of the project, and provided overall direction, mentoring, and manuscript editing. JL provided technical design advice, code, and contributed to the text. CF provided project management, trial curation, and contributed to the text. ZZ was the primary software developer and dev ops engineer. PK, BV, MT, and TM contributed to the code and infrastructure. Additionally, PK was the data acquisition lead. SB, AA, KD, BB, and GS aided in deployment, testing, and facilitated early adoption of the platform. JM, LS, LM, NL, BJ, BR, CS, and MH acted as a steering committee and addressed regulatory issues. All authors read and approved the final version of the manuscript.

## Acknowledgements

The authors would like to thank Laura Kleiman for editing the manuscript, and the Early Drug Development Center at DFCI, especially Khanh Do, for their persistent use of MatchMiner to match patients to trials.

## Additional Files

Supplementary table 1 — Genes and their usage in clinical trial eligibility.

All genes listed as inclusion or exclusion criteria for the 178 curated clinical trials at DFCI. Genes are sorted by clinical trial count.

